# A positively selected, common, missense variant in *FBN1* confers a 2.2 centimeter reduction of height in the Peruvian population

**DOI:** 10.1101/561241

**Authors:** Samira Asgari, Yang Luo, Gillian M. Belbin, Eric Bartell, Roger Calderon, Kamil Slowikowski, Carmen Contreras, Rosa Yataco, Jerome T. Galea, Judith Jimenez, Julia M. Coit, Chandel Farroñay, Rosalynn M. Nazarian, Timothy D. O’Connor, Harry C. Dietz, Joel Hirschhorn, Heinner Guio, Leonid Lecca, Eimear E. Kenny, Esther Freeman, Megan B. Murray, Soumya Raychaudhuri

## Abstract

Peruvians are among the shortest people in the world. To understand the genetic basis of short stature in Peru, we examined an ethnically diverse group of Peruvians and identified a novel, population-specific, missense variant in *FBN1* (E1297G) that is significantly associated with lower height in the Peruvian population. Each copy of the minor allele (frequency = 4.7%) reduces height by 2.2 cm (4.4 cm in homozygous individuals). This is the largest effect size known for a common height-associated variant. This variant shows strong evidence of positive selection within the Peruvian population and is significantly more frequent in Native American populations from coastal regions of Peru compared to populations from the Andes or the Amazon, suggesting that short stature in Peruvians is the result of adaptation to the coastal environment.

**One Sentence Summary:** A mutation found in Peruvians has the largest known effect on height for a common variant. This variant is specific to Native American ancestry.

## Main Text

With average male and female heights of 165.3 cm and 152.9 cm respectively, Peruvians are among the shortest people in the world (1). The genetic makeup of current Peruvians is shaped by extensive admixture between Native American residents of Peru and the incoming Europeans, Africans, and Asians who arrived in Peru since 18 century (2, 3). In a 2014 genetic study of individuals from South and Latin America, Ruiz-Linares *et al.* reported that Native American ancestry is correlated with lower height (4). However, as the authors acknowledged, this association may have been the result of confounding socioeconomic status or environmental factors related to indigeneity, which could not be captured by socioeconomic variables included in this study (education and wealth). Even in the case where the observed correlation between Native American ancestry and height were driven at least in part by genetic factors, it is unclear what genes or adaptive processes might have driven this effect.

In order to define distinctive genetic factors contributing to height in Peruvians, we performed the first large-scale genetic study of height in Peru (**Methods**). We obtained height and genotyping data from 1,795 males (57%) and 1,339 females (43%) in 1,947 households (**Fig S1**) and inferred the level of Native American ancestry in each individual **(Methods, Fig S2, FigS3)**. We observed a negative correlation between height and Native American ancestry proportion (Pearson correlation coefficient (*r*) = −0.3, p-value=9.3×10^−58^, **Fig. 1A**, **Fig. S4A**). Native American ancestry remained significantly associated with lower height after including age, gender, African, and Asian ancestry proportions and a random household effect in the model, as a proxy for unmeasured environmental factors (p-value =7.2×10^−43^, effect size = −14.7 cm, SE = 1.1, **Fig. 1B**, **Table S1, Table S2, Fig. S4B**).

**Fig. 1:**
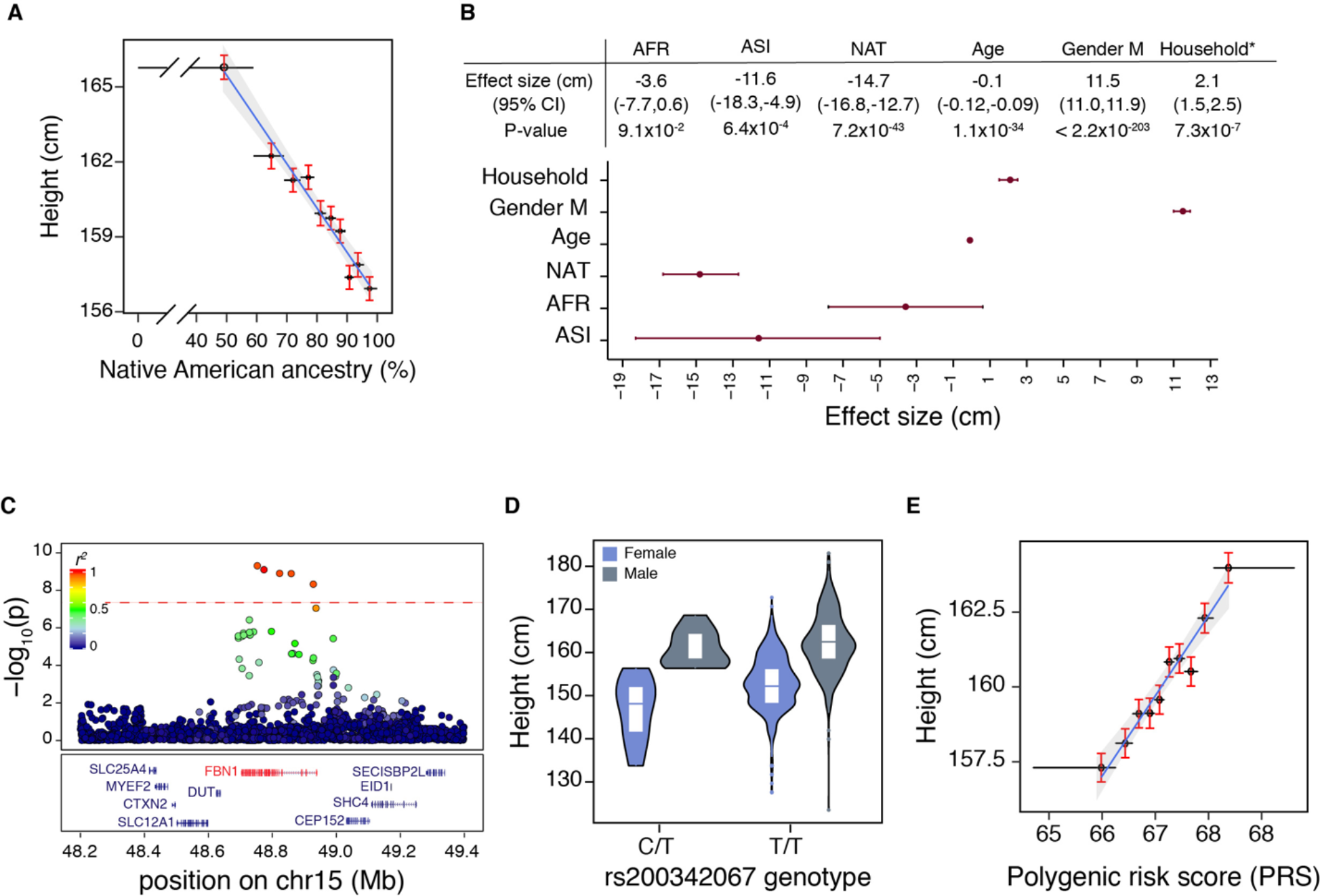
Genetic architecture of height in the Peruvian population. **A)** Height is negatively correlated with the proportion of Native American ancestry (*r* = −0.98, p-value=3.7×10^−7^). Each point represents the median for a decile of Native American ancestry (x-axis) and the average height for that decile (y-axis). Error bars show range (x-axis) and standard error (SE, y-axis). **B)** Increased Native American ancestry is significantly associated with lower height after accounting for age, gender, African and Asian ancestry proportions, household as a proxy for socioeconomic factors, and cryptic genetic relatedness. *Household effect size is calculated as the standard deviation (SD) in the model’s intercept. **C)** Locus-specific Manhattan plot of *-log_10_* transformed p-values from GWAS of height in the Peruvian population. One locus on chromosome 15 passed the genome-wide significance threshold (p-value < 5×10^−8^, N = 3,134). This locus includes five SNPs in *FBN1*, including one population-specific missense variant (rs200342067, MAF = 4.7%, p-value = 8.0×10^−10^, effect size = −2.2 cm, SE = 0.4). **D)** rs200342067 showed a similar effect size in an independent cohort of individuals with Mexican, South American, and Central American ancestry (N = 1,935, MAF = 0.7%, p-value = 6.3×10^−4^, effect size = −2.2 cm, SE = 0.6) **E)** Height is positively correlated with polygenic risk scores (PRS) (*r* = 0.97, p-value = 2.3×10^−6^). Each point represents the median for a PRS decile (x-axis) and the average height for that decile (y-axis). Error bars show range (x-axis) and SE (y-axis).

To test whether specific variants derive this effect, we performed a genome-wide association study (GWAS). One locus on chromosome 15 (15q21.1) passed the genome-wide significance threshold (p-value < 5×10^−8^, **Fig. S5A**). This locus overlaps the coding sequence of *FBN1* (15q15-21.1) and includes five single nucleotide polymorphisms (SNPs). One SNP, rs200342067 (minor allele frequency (MAF) = 4.7%, p-value = 8.0×10^−10^, effect size = −2.2 cm, SE = 0.4), is a missense variant while the other four are intronic variants (**Fig. 1C**, **Table S3, Table S4**). In the 1000 Genomes Project (5), rs200342067 is specific to the Mexican (MAF = 0.8%) and Peruvian (MAF = 4.1%) populations and is not found in the European, African, and Asian populations. We examined the association of this variant with height in an independent cohort of 1,935 individuals with Mexican, Central American, and South American ancestry from the Bio*Me* Biobank at the Icahn School of Medicine at Mount Sinai in New York City (**Table S5**). We observed an identical effect size (MAF = 0.7%, p-value = 6.3×10^−4^, effect size = −2.2 cm, SE = 0.6, **Fig. 1D**), independently replicating the association, and confirming that this association is not driven by technical bias or other non-genetic factors. We did not find any additional association in a gene-based analysis of rare (MAF < 1%) variants or a gene-based meta-analysis of common variants (**Methods, Fig. S5B-C**).

Previous large-scale height GWAS, done predominantly in Europeans, have identified 3,290 independent common height-associated variants (6). To assess the predictive power of these European-biased variants in the Peruvian population we generated polygenic risk scores (PRS) based on the reported effect sizes of 2,993 common height-associated variants that were present in our cohort (**Methods, Fig. S6A**). Higher PRS bins were associated with increased height (*r* = 0.2, p-value = 2.7×10^−34^, **Fig. 1E**, **Fig. S6B**). The estimated genetic heritability 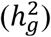 of height was similar for Peruvians (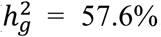, **Methods**) and Europeans 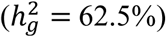 (7); however, previously identified height-associated variants explained only 6.1% of height phenotypic variance in our cohort compared to 24.6% in the original European cohort (**Methods, Fig. S6C-D**), suggesting that either different variants are responsible for the height variance in the Peruvian population or the lead European variants do not tag the same causal variants in the Peruvian population. This observation is in line with a number of recent reports (8–10) showing the lower predictive power of PRS calculated based on European GWAS in non-European populations as a result of differences in demographic history and linkage disequilibrium (LD) patterns.

Of previously identified common height-associated variants, 99% have effects less than 5 mm per allele (**Fig. S6E**). In contrast, rs200342067 in heterozygous individuals reduces height by 2.2 cm (4.4 cm in homozygous individuals) and explains 0.9% of height phenotypic variance in our cohort (**Fig. S7**). This effect size is comparable to the effect sizes seen for extremely rare height-associated variants that are believed to be under purifying selection (**Fig. 2A**) (6, 11). In fact, almost all missense variants in *FBN1* are under purifying selection, causing this gene to have a significantly lower burden of missense variants compared to what is expected by chance (z-score = 5.53, p-value = 3.2×10^−8^, Exome Aggregation Consortium (ExAC), N = 60,706) (12). Given the large effect size and high MAF of rs200342067, we hypothesized that this variant is under positive selection in the Peruvian population.

**Fig. 2:**
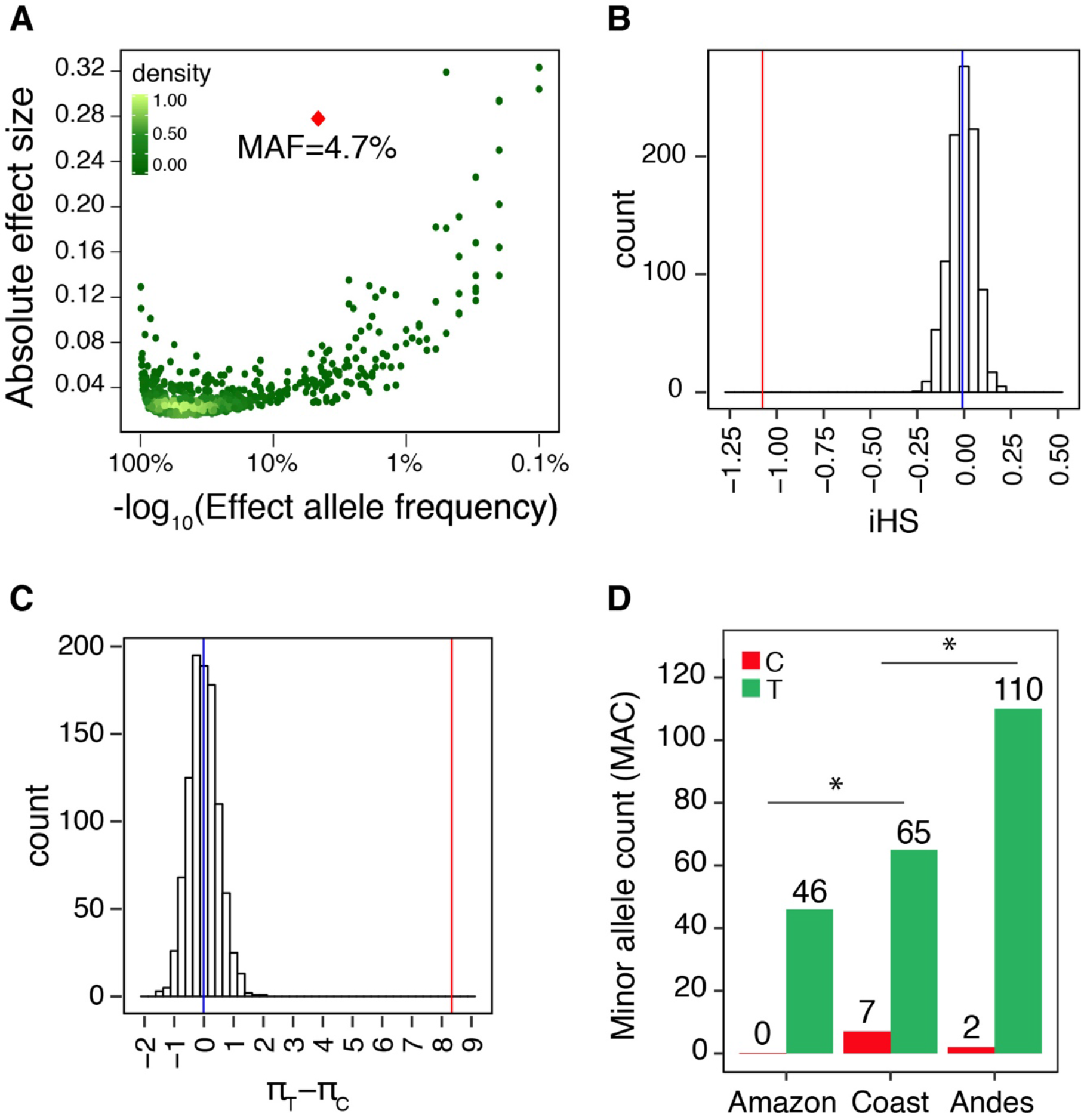
rs200342067 is positively selected in the Peruvian population. **A)** Effect sizes and allele frequencies of 3,290 previously identified height-associated variants in the European population (6) (green dots) compared with the effect size and allele frequency of rs200342067 (red diamond) from this study (MAF = 4.7% in the Peruvian population and zero in the Europeans). To compare different studies, the effect size is shown as the absolute effect size on scaled height values. Haplotypes carrying the C allele at rs200342067 **B)** are significantly longer (observed iHS = −1.1, null distribution range = −0.25 to 0.26) and **C)** have significantly lower nucleotide diversity (observed π_C_-π_T_ = −8.5, null distribution range = −1.7 to 1.4) compared to the haplotypes carrying the T allele. **D)** The rs200342067 variant was significantly more frequent in the populations from the coastal regions compared to the populations from the Andes or the Amazon (two-sided Fisher’s exact test p-value = 0.01, 0.02, and 0.005 for coast versus Amazon, coast versus Andes, and coast versus Andes and Amazon).

We used Integrated Haplotype Score (iHS) (13) and pairwise nucleotide diversity (π) (14) statistics to search for signals of positive selection in a 1Mb window around rs200342067 within the Peruvian population. We restricted these analyses to haplotypes in which the local ancestry of the core SNP (e.g. rs200342067) was inferred as Native American (N = 271 with C and 4,720 with T at the core SNP, **Methods, Fig. S8**). To generate an empirical null distribution, we randomly assigned C (derived) and T (ancestral) alleles to rs200342067 1000 times and calculated iHS and π in each round (**Methods).** Haplotypes carrying the C allele were significantly longer (observed iHS = −1.1, permutation mean = 0 and range from −0.27 to 0.19, **Fig. 2B**) and had a significantly lower number of pairwise sequence differences (observed π_T_-π_C_ = −8.3, permutation mean = −0.01 and range from −1.5 to 1.9) compared to the haplotypes carrying the T allele (**Fig. 2C**). Haplotypes carrying the C allele also showed a slower decay of homozygosity (EHH), from the core SNP (**Methods, Fig. S9**) (15).

Since adaptation to local environment can drive dramatic allele frequency shifts, we decided to compare the frequency of rs200342067 among Peruvian populations from different geographical regions in Peru (N = 115) (3). The rs200342067 variant was significantly more frequent in the individuals from coastal regions compared to the individuals from the Andes or the Amazon (MAF = 9.7%, 1.7%, and 0 for Coast, Andes and Amazon respectively, coast vs. non-coast two-sided Fisher’s exact test p-value = 0.005, **Fig. 2D**, **Table S6**). Among the coastal populations the Moche population, a native population in the North coast of Peru, had an especially high frequency of rs200342067 (N = 21, minor allele count = 4, MAF= 9.5%). Notably, the average height of Moche people is far below the average height in Peru (158 cm and 147 cm for Moche male and female (16) versus 164 cm and 152 cm for the average Peruvian male and female in the same year (1)), suggesting that rs200342067 might have been selected in the Peruvian population as a result of adaptation to the coastal environment.

The rs200342067 variant changes the conserved T (ancestral) allele to a C (derived) allele in *FBN1* exon 31 (g.48773926T>C, **Fig. S10, Fig. S11A-B**). This change substitutes a large, negatively charged glutamic acid for a glycine, the smallest amino acid with no side chain in fiibrillin-1 (E1297G), the protein encoded by *FBN1*. Fibrillin-1 is an extracellular matrix (ECM) glycoprotein that serves as a structural backbone of force-bearing microfibrils in elastic and non-elastic tissues (17) and is also involved in tissue development, homeostasis, and repair by interacting with different growth factors including transforming growth factor (TGFβ) (17). E1297G is located in fibrillin-1 calcium binding epidermal growth factor domain 17 (cbEGF-domain 17), between a conserved cysteine (C1296) involved in forming a disulfide bond with C1284, and a conserved asparagine (N1298) involved in calcium binding (18) (**Fig. S11C-D**).

While the clinical significance of fibrillin-1 E1297G is not known, other missense mutations in fibrillin-1 are known to cause nine different Mendelian diseases (**Table S7**), all of which are dominantly inherited (19). Most of fibrillin-1-realted Mendelian diseases include skeletal anomalies and changes in skin elasticity among other symptoms (19). To investigate the possible clinical consequences of fibrillin-1 E1297G we performed clinical exams on 11 individuals from our cohort: 2 homozygous (C/C) cases, 2 heterozygous (C/T) cases, and 7 matched controls with reference (T/T) genotype (**Table S8, Methods**). Musculoskeletal examination on these individuals did not show any obvious difference in the range of motion of knees, hips, wrists, and proximal interphalangeal and metacarpophalangeal joints of the second and third digits. One C/C genotype individual (individual-1, **Table S8**) had notably thicker skin upon a total body skin examination, and appeared much older than the stated age, while the other C/C genotype individual had no clinically abnormal cutaneous findings. None of the C/T or T/T individuals had an abnormal skin exam.

While all the reported fibrillin-1 mutations causing short stature phenotypes occur in the TGFβ-binding domains (**Fig. S11C**), mutations in the cbEGF-domains of fibrillin-1 (**Fig. S11C**), especially the ones affecting cysteine residues or residues involved in calcium binding, lead to Marfan or Marfan-like syndromes (20). Notably, missense mutations in cbEGF-domains 11 to 18 of fibrillin-1 (a.k.a. neonatal region, **Fig. S11C**) are previously associated with severe neonatal forms of Marfan syndrome, mortality within the first two years of postnatal life, and poor disease prognosis in adults (19, 21). To our knowledge, E1297G is the first report of a mutation in the fibrillin-1’s neonatal region that leads to short stature, a manifestation opposite to the higher stature common in Marfan syndrome.

Common variants with large effect sizes on height might increase in frequency in population as a result of positive selection. A study of height in Sardinian islanders found an intronic variant in *KCNQ1*, which encodes a voltage-gated potassium channel, that reduces height by an average of 1.8 cm (rs150199504, MAF = 7.7%, MAF in Central European population (CEU) = 0.67%); they suggested that this variant is positively selected in Sardinians as a result of adaptation to the island environment (22). A study of signatures of genetic adaptation in Greenland Inuits found an intronic variant in *FADS3*, a gene involved in fatty acid metabolism, that reduces height by 1.9 cm possibly due to the influence of fatty acid composition on the regulation of growth hormones (rs7115739, derived allele frequency (DAF) = 62.7%-81.9%, DAF in CEU = 2.9%-3.6%); they suggested that this variant is positively selected in Greenland Inuits as a result of adaptation to the their fat-rich diet (23). Similarly, it is plausible that short stature is the result of adaptation to the coastal environment. It is also possible that other *FBN1*-related traits like changes in cardiovascular system performance have offered an evolutionary advantage in the Native American ancestors of Peruvians. Understanding the exact adaptive processes that could have caused the selection of rs200342067 in the Peruvian population is a challenging task and requires further investigation.

Besides its implications in medical and population genetics, this study highlights the importance of large-scale genetic studies in underrepresented and founder populations. Our findings show that genetic studies in novel populations can uncover novel trait-associated variants of large effects in functionally relevant genes. Similar studies in diverse populations are required to capture the extent of human genetic diversity and to expand the benefits of genetic research to all human populations.

## Acknowledgments

The study was supported by the National Institutes of Health (NIH) TB Research Unit Network, Grants U19-AI111224-01 and U01-HG009088. The content is solely the responsibility of the authors and does not necessarily represent the official views of the NIH. S. Asgari was supported by the Swiss National Science Foundation (SNSF) Early postdoc mobility fellowship P2ELP3_172101;

## Author contributions

S.R. and M.M. designed the study. S.A. analysed and interpreted the data. S.A. and S.R. drafted the manuscript. Y.L., G.B. E.K., J.H., E.B., K.S., H.G., and T.O., performed statistical analysis. M.M., L.L., J.C., C.C., R.Y., J.G., J.J., J.C., and C.D. recruited patients and obtained samples for this study. S.R., E.F., H.D., and R.N. conducted clinical assessment.;

## Data and materials availability

Genotyping data will be made available through dbGAP.

## Supplementary Materials

**Figure S1: Cohort’s demographic information. A)** Density plot of height for all the Peruvian males (N = 1,795 (57%)) and females (N = 1,339 (43%)) included in this study after quality control (e.g. after removing low `quality samples, individuals below 18 years old and height outliers (土 3 x standard deviations (SD) from the mean`). Males were significantly taller than females (Male mean = 165.2 cm (SD = 6.7), Female mean = 153.4 cm (SD = 6.4), p-value < 2.2×10^−308^). **B**) Age was not significantly different between males and females (t-test p-value = 0.09).

**Figure S2: Principal component analysis (PCA).** PCA analysis of genotyping data from Peruvians included in this study merged with the data from continental populations from the 1000 Genomes Project phase 3 (N = 3469) (5, 24) as well as the data from Siberian and Native American populations from Reich et *al*. 2012 *Nature* study (25) (N = 738) as reference panel (number of variants = 34,936, MAF > 1%, genotype missingness < 5%). In order to better visualize the relative position of reference populations we plotted the data **A)** without and **B)** with the Peruvians from this study (N = 3,134). Each individual is represented as a dot. Populations are colored based on their continental origin for the 1000 Genomes Project data and based on assignment to Native American or Siberian tribes for the Reich data (AFR: Africa, AMR: South America, EAS: East Asia, SAS: South Asia, EUR: Europe, SIB: Siberian, NAT: Native American).

**Figure S3: Global ancestry analysis using ADMIXTURE (K=4)**. We observed varying levels of European, African, and Asian admixture in the Peruvian population with a median proportion of Native American, European, African, and Asian ancestry per individual of 0.83 (Interquartile range (IQR) = 0.72-.91), 0.14 (0.08-0.21), 0.01 (0.003-0.03), and 0.003 (10^−5^-0.01) respectively. Each individual is represented as a thin vertical line, each color corresponds to the genomic proportion of a given cluster in that individual’s genome. The ADMIXTURE with K=4 analysis is done using all populations in 1000 Genomes Project phase 3 (5, 24) and Siberian and Native American populations from the Reich et *al*. 2012 *Nature* study (25). AFR: African ancestry includes: Yoruba in Ibadan, Nigeria, Luhya in Webuye, Kenya, Gambian in Western Divisions in the Gambia, Mende in Sierra Leone, Esan in Nigeria, Americans of African Ancestry in SW USA; EUR: European ancestry, includes: Central European, Utah Residents (CEPH) with Northern and Western European Ancestry, Toscani in Italy, Finnish in Finland, British in England and Scotland, Iberian Population in Spain; EAS: East Asian, includes: Han Chinese in Beijing, China, Japanese in Tokyo, Japan, Southern Han Chinese, Chinese Dai in Xishuangbanna, China, Kinh in Ho Chi Minh City, Vietnam; SAS: South Asian, includes: Gujarati Indian from Houston, Texas, Punjabi from Lahore, Pakistan, Bengali from Bangladesh, Sri Lankan Tamil from the UK, Indian Telugu from the UK; PUR: Puerto Ricans from Puerto Rico; CLM: Colombian from Medellin, Colombia; MXL: Mexicans from Los Angeles, California; PEL: Peruvians from Lima, Peru. Altic: Altaic language family, includes: Yakut, Buryat, Evenki, Tuvinians, Altaian, Mongolian, Dolgan. North Amerind: Northern Amerindian language family, includes: Maya, Mixe, Kaqchikel, Algonquin, Ojibwa, and Cree. Central Amerind: Central Amerindian language family, includes: Pima, Chorotega, Tepehuano, Zapotec, Mixtec, and Yaqui. Andean: Andean language family, includes: Quechua, Aymara, Inga, Chilote, Diaguita, Chono, Hulliche, and Yaghan. For a full list of all populations in all language groups please see the Reich et *al*. 2012 *Nature* study (25).

**Figure S4: The effect of Native American ancestry proportion on height. A)** Greater Native American ancestry proportion is associated with lower height (N=3,134, *r* = −0.3, p-value = 9.3×10^−58^). The x-axis represents Native American ancestry proportion from ADMIXTURE analysis at K = 4 clusters. The y-axis represents height (cm). **B)** Height was randomly reassigned to individuals within each household, and the effect size of Native American ancestry on height was recalculated to derive an empirical null distribution of effect sizes. None of the permutations resulted in a greater effect size than that of the original data (permutation effect size ranging from −5.6 cm to 5.8 cm, permutation mean effect size = 0 cm, observed effect size = −14.7 cm)

**Figure S5: Manhattan and quantile-quantile (QQ) plots. A)** Single variant association analysis using GEMMA (26), the dotted red line corresponds to the genome-wide significance threshold of 5×10^−8^ for single variant association testing. Five SNPs passed the genome-wide significance threshold. **B)** Rare (MAF < 1%) variants gene-based analysis using SKAT (27) the dotted red line corresponds to the genome-wide significance threshold of 2×10^−6^ for 25,000 tested genes. No SNPs reached the genome-wide significance threshold. **C)** gene-based meta-analysis of common variants using GCTA fastBAT (28) the dotted red line corresponds to the genome-wide significance threshold of 2×10^−6^ for 25,000 tested genes. No SNPs reached the genome-wide significance threshold.

**Figure S6: Polygenic risk score (PRS) analysis.** We used effect sizes from 2,993 common height-associated variants from the Yengo et al 2018 meta-analysis (N ∼ 700,000 European individuals) (6) that were present in our cohort (N = 3,134 Peruvian individuals) to derive the PRS. **A)** Out of 2,993 variants, 1,519 (51%) showed directionally consistent effects, and 199 (7%) had p-value < 0.05 in our Peruvian GWAS. **B)** Higher PRS values are associated with increased height (*r* = 0.2, p-value = 1.7×10^−34^)**. C)** Histogram showing the PRS distribution. **D)** Previously identified height-associated variants explained only 6.1% of height phenotypic variance in our cohort (*r* = 0.061, p-value = 6.8×10^−45^), x-axis: PRS, y-axis: height residuals after adjustments for age and gender as fixed effects and a GRM as random effect. **E)** The majority (99%) of previously identified common height-associated variants (N = 3,290) have effects less than 5 mm per allele (dashed red line: cutoff corresponding to 5 mm effect size, smaller plot shows the zoomed in tail of the main plot).

**Figure S7: Effect size of rs200342067 on height in the Peruvian population**. rs200342067 in heterozygous individuals reduces height by 2.2 cm (4.4 cm in homozygous individuals, including 11 individuals with C/C genotype, 275 C/T genotype, and 2,848 T/T genotype) and could explain 0.9% of height phenotypic variance in our cohort (N = 3,143). x-axis: rs200342067 genotype, y-axis: height residuals after adjustments for age and gender as fixed effects and a GRM as random effect.

**Figure S8: Local ancestry inference at the rs20034206 locus.** To test for positive selection at the rs20034206 locus, we restricted the analysis to haplotypes in which the local ancestries of both C and T alleles were inferred to be Native American. **A)** Local ancestry inference results for A) rs20034206 C allele, and **B)** rs20034206 T allele.

**Figure S9: Extended haplotype homozygosity (EHH) for rs20034206, C and T alleles.** Haplotypes carrying the C allele show a slower decay of homozygosity compared to the haplotypes carrying the T allele. Analysis is restricted to haplotypes in which rs20034206 is inferred as Native American.

**Figure S10: Multiple sequence alignment around rs20034206 in 37 eutherian mammals**. rs200342067 changes a conserved T allele (ancestral, shown in red) to a C allele (derived). Sequence alignments were obtained from Ensembl GRCh37 release 95.

**Figure S11: Genomic context of rs200342067 (E1297G). A)** Schematic representation of *FBN1*, exons are shown as black bars. Exon 31 (ENSE00001753582) is shown in red. B) *FBN1* exon 31 sequence and PhyloP per-nucleotide conservation score based on multiple alignment of 100 vertebrate species (obtained from UCSC genome browser GRCh37 assembly, conservation track). The T>C change due to rs200342067 occurs in a conserved nucleotide. **C)** Schematic representation of Fibrillin-1 (ENST00000316623.5). Fibrillin-1 consists of: N and C terminal (black rectangles), EGF-like domains (stripped rectangles), hybrid domains (black pentagons), TGFβ-binding domains (gray ovals), a proline-rich domain (white hexagon), and 43 calcium binding cbEGF-like domains (white rectangles). cbEGF-domain 17, the domain affected by rs200342067 (p.Glu1297Gly), is shown in red. p.Glu1297Gly is located between a conserved cysteine (p.Cys1296) involved in forming a disulfide bond with p.Cys1284 and a conserved asparagine (p.Asp1298) involved in calcium binding. **D)** Fibrillin-1 cbEGF-domain 17 sequence and 3D structure of cbEGF-domains 17 and 18 (the 3D structure was obtained based homology with fibrillin-1 cbEGF-domains 12 and 13, 1LMJ (18), in the Protein Data Bank). rs200342067 changes glutamic acid, a large amino acid with a negatively charged side chain, to glycine, the smallest amino acid with no side chain (shown in red). The side chains are shown for rs200342067 (red spheres), the calcium-interacting residues (beige sticks), and the cysteine residues involved in disulfide bonds (yellow sticks). Calcium ion is shown in green.

**Table S1: Base Model parameters.** Native American ancestry is significantly associated with lower height after accounting for age, gender, African and Asian ancestry proportions, and a genetic relatedness matrix (GRM) to account for population structure and genetic relatedness. ASI: Asian, AFR: African, EUR: European, NAT: Native American.

**Table S2: Base model plus a household random effect.** Native American ancestry remained significantly associated with lower height after we included a random household effect as a proxy for socioeconomic and environmental factors.

**Table S3: Association signal at 15q15-21.1**. This locus overlaps the coding sequence of *FBN1* and includes five single nucleotide polymorphisms (SNPs), which are in high LD.

**Table S4: Inclusion of other covariates in rs200342067 association testing.** Inclusion of principal components (PCs), socioeconomic status (SES), or ancestry proportions (ASI: Asian, AFR: African, EUR: European) did not change the association effect size or strength.

**Table S5: rs200342067, C genotype carriers in *BioMe* cohort**. Individuals are stratified by country of origin. No homozygous individual (C/C) was observed in *BioMe.* `For the replication analysis, we restricted the age to ≥ 18 and ≤ 80 for females, and ≥ 22 and` ≤ 80 for males.

**Table S6: Comparison of rs200342067 minor allele count between populations from different geographical regions in Peru.** rs200342067 was significantly more frequent in Coastal populations than in populations from the Andes and the Amazon.

**Table S7: Disease, phenotypes, and traits caused by mutations in FBN1.**

**Table S8: Demographic information of clinical examination participants.** Skin biopsies were obtained from 11 including: 2 with C/C, 2 with C/T, and 7 with T/T genotypes at rs200342067.

