## supplementary material for "A positively selected, common, missense variant in *FBN1* confers a 2.2 centimeter reduction of height in the Peruvian population"

**This PDF file includes:**

Materials and Methods

Supplementary text

Figures S1-S11

Tables S1-S8

References (*1-37*)

### Material and methods

#### Study design & participants

As described previously (*1*), the data was obtained following institutional IRB guidelines and with informed consent from participants. We collected genotyping data for 4,002 individuals from 1,769 households in Lima, Peru, using a customized Affymetrix Axiom array as described previously (*1*). In brief, we designed a ~720K marker array based on exome-sequencing data from 116 Peruvians in order to optimize for population-specific rare and coding variants. Quality control on the genotypes, phasing and imputation was performed as described previously using GRCh37 as the reference genome (*1*).

#### Phenotype

Height in centimeters, gender, age, socioeconomic status, and individuals' TB status were collected. We excluded 846 individuals from the analysis: individuals below 19 years of age, individuals without height measurement, and individuals with a measured height more than ± three standard deviations (3*SD) away from the population average.

#### Kinship estimation

Many kinship estimation methods perform under the assumption of sampling from a single population with no underlying ancestral diversity. Kinships estimates are inflated when this assumption is violated (*2*). In the presence of population structure and admixture, methods that replace population allele frequencies with ancestry-specific allele frequencies are preferred (*2*). We used the GENESIS R package (version 2.6.1) to estimate the kinship coefficients between individuals. The package is based on a method called PC-Relate (*3*), which uses ancestry representative PCs to correct for population structure. Individuals were considered unrelated if their estimated kinship coefficients were ≤ 0.0625, corresponding to second degree genetic relatedness or closer. 476 individuals had kinship coefficients > 0.0625.

#### Genetic relatedness matrix (GRM)

To avoid spurious association results it is important to account for both recent genetic relatedness, such as family structure, and more distant genetic relatedness, such as population structure. To this end, we used GEMMA (*4*) (version 0.96), with default options, to generate a GRM after removing rare variants (MAF ≤ 1%), regions with known long-range linkage disequilibrium (LD) (*5*), and variants in high LD ($r^2 > 0.2$ in a window of 50kb and a sliding window of 5kb). We used PLINK (version 1.90b3w) for pruning the genotypes.

#### Principal Component Analysis (PCA)

We merged our genotype data with data from the continental populations of phase 3 of the 1000 Genomes Project (*6*, *7*) and genotype data from Siberian and Native American populations from the Reich et *al*. 2012 *Nature* study (*8*) by matching on chromosome, position, reference, and alternate alleles. After merging the datasets, variants with an overall MAF < 1% were excluded. We used GCTA (*9*) (version 1.26.0) to perform PCA. We used PLINK (version 1.90b3w) (*10*) for LD pruning, merging, and quality control. The merged dataset included 34,936 variants.

#### Global ancestry inference

We used ADMIXTURE (*11*) (version 1.3) at K = 4 clusters, for global ancestry inference. The choice of four ancestral populations for ADMIXTURE analysis was based on Peru's demographic history and previous studies of Peruvian population structure (*12–14*). We used the merged dataset described above as input for the ADMIXTURE analysis. We used PLINK (version 1.90b3w) (*10*) to exclude variants with genotyping missingness rate > 5% and to perform LD pruning by removing the markers with $r^2 > 0.1$ with any other marker within a sliding window of 50 markers per window and an offset of 10 markers.

**Local ancestry inference**

We phased our data using SHAPEIT2 (*15*) (version v2.r837) and converted all files to RFMix format using publicly available scripts (*16*). For local ancestry inference we included the following populations from the 1000 genomes project (*17*) as reference populations: YRI for African ancestry, CEU for European ancestry, and PEL with inferred Native American ancestry > 0.85 based on ADMIXTURE at K = 4 clusters analysis, as a proxy for Native American ancestry. We inferred the local ancestry on the phased haplotypes using RFMix (*18*) (version 1.5.4). We ran RFMix with the following flags "-n 10 -w 0.1 -e 1 --skip-check-input-format --num-threads 10 --use-reference-panels-in-EM --forward-backward".

**Correlation between global ancestry proportions and height**

We used the R package lme4qtl (*19*), a linear mixed model framework, to measure the correlation between global ancestry proportions and height. We included the following covariates in the base model: age, gender, African and Asian ancestry proportions, as well as a GRM to account for population structure and genetic relatedness. We repeated this analysis after adding of a random effect to account for individual's household. To ensure adequate control for environmental factors, we randomly assigned height to individuals within each household 10,000 times and recalculated the Native American ancestry effect size using the base model to generate an empirical null distribution. We compared the null distribution with the observed Native American ancestry effect size from the original data to generate an empirical permutation p-value.

**Common variants association analysis**

We limited the analysis to variants having an overall MAF>1%, Hardy-Weinberg p-value (HWE-P) $>10^{-5}$, and an overall INFO score>0.3, using PLINK version (1.90b3w) (*10*). We split the multi-allelic variants into multiple variants, creating a single variant for each alternate allele. We used GEMMA (*4*) (version 0.96) to perform the single variant genome-wide association analysis, with age, gender, and GRM as covariates. We repeated the association for chromosome 15 by adding one or more of the following covariates: 10 PCs, 20 PCs, socioeconomic status, African global ancestry proportion, Asian global ancestry proportion, and European global ancestry proportion. For the replication analysis we used genotyping data from 1,935 individuals with Mexican, Central American, and South American ancestry from the Bio*Me* Biobank at the Icahn School of Medicine at Mount Sinai in New York City using the lm() function in R (v.3.2.0). We restricted the age to $\geq$ 18 and $\leq$ 80 for females, and $\geq$ 22 and $\leq$ 80 for males. Height in centimeters was used as the outcome variable with rs200342067 genotype status as the primary predictor variable and gender as covariate (with N = 25 carriers of the "C" allele in total).

**Heritability analysis**

We used GREML analysis in GCTA (*20*) (version 1.26.0) to calculate the amount of variance in height explained by all common variants (MAF > 1%). We included 423,108 variants from 2,667 unrelated individuals in this analysis with age, gender, and the first 10 PCs as covariates in the analysis.

**Polygenic risk score (PRS) analysis**

We constructed polygenic risk scores (PRSs) for each individual using height-increasing effect sizes from 2,993 previously published independent height-associated variants (*21*) as follow:

$$PRS_j = \sum_{i=1}^{m} n_{ij} * \beta_i$$

Where $\beta_i$ is the reported effect size for variant *i*, $n_{ij}$ is the allele count of variant *i* in individual *j* and *m* is the total number of variants used in the construction of the PRS.

**Gene-based association analysis**

We used SKAT (*22*) (version 1.3.2.1) for gene-based association testing of rare (MAF ″ 1%) variants. We restricted the analysis to variants with INFO score > 0.3, HWE < $10^{-5}$. Null distributions were generated using SKAT_NULL_emmaX, which incorporates kinship structure in the calculation of SKAT parameters and residuals. Age and gender were included as covariates. Statistical significance threshold was set at $p < 2.5\times10^{-6}$ which is the Bonferroni correction threshold for 20,000 protein coding genes. For common variants (MAF > 1%) we used fastBAT analysis in GCTA (*23*) to perform gene-based association testing using GWAS summary statistics.

**Test of positive selection**

We used selscan (*24*) (version 1.2.0a) to calculate extended haplotype homozygosity (EHH) (*25*), integrated haplotype score (iHS) and the mean pairwise number of nucleotide differences (nucleotide diversity, $\pi$) (*26*) on phased genotypes with MAF > 1%. We restricted the analysis to haplotypes in which the inferred ancestry of rs200342067 was Native American (see RFMix methods). We calculated iHS and $\pi$ in a 1Mb window around rs200342067. Both EHH and iHS are statistics based on the increased LD around the positively selected allele compared to the non-selected allele.. Negative iHS values indicate that haplotypes surrounding the derived allele are longer compared to the haplotypes surrounding the ancestral allele, implying positive selection at the derived allele (*27*). Positive selection reduces genetic diversity at the site of selection (*28*); the $\pi$ metric assess positive selection by measuring the average number of pairwise sequence differences between two randomly selected haplotypes and is expected to be lower for haplotypes surrounding the positively selected allele. To test the significance of our results, we generated an empirical null distribution by randomly assigning C (derived, minor) and T (ancestral, major) alleles to rs200342067 at each haplotype, while keeping the total number of C and T haplotypes identical to the original data, 1000 times and calculating iHS and $\pi$ in each round. We compared the null distributions with the observed iHS and $\pi$ values from the original data to generate an empirical p-value. Minor allele counts for rs200342067 in populations from different geographical regions in Peru were obtained from the study by Harris et al. (*14*). We used Fisher's exact test in R (version 3.4) to test the significance of the observed differences in minor allele counts.

**FBN1 cbEGF-domain 17, 3D structure**

The 3D structure was obtained based homology with fibrillin-1 cbEGF-domains 12 and 13, 1LMJ, in the Protein Data Bank (PDB) (*29*).

**Clinical examination**

Clinical examination and collection of skin biopsies from study participants was approved by the local Institutional Review Board (IRB). Individuals with T/T genotype (controls) were matched with cases (individuals with C/C and C/T genotypes) for gender, age$\pm$5 years, Native American ancestry proportion$\pm$0.05, and European ancestry proportion$\pm$0.05. A board-certified rheumatologist performed a musculoskeletal exam and history, including a detailed musculoskeletal history with review of systems, past medical history, medication history, social history, and family history; vital signs; range of motion on knees, wrists, elbows, index fingers, middle fingers, and hips; joint exam of hands for bony changes, synovitis or other abnormalities; joint exam of knees, feet, and spine for instability, bony changes, inflammation or other abnormalities. A board-certified dermatologist performed a standardized total body skin exam. This includes examination of the skin of the face, eyelids, ears, scalp, neck, chest, axillae, abdomen, back, buttocks, genitalia, upper extremities, lower extremities, hands, feet, digits, nails, lips, mouth, mucosae, and lymph nodes.

**Supplementary text**

**Permutation analysis to test the association between Native American ancestry and height**

We observed a negative correlation between height and Native American ancestry proportion after correcting for age, gender, African, and Asian ancestry proportions and a random household effect in the model, as a proxy for unmeasured environmental factors (p-value =$7.2 \times 10^{-43}$, effect size = -14.7 cm, SE = 1.1). To ensure adequate control for environmental factors, we also performed a stringent permutation analysis "within each household". We randomly reassigned heights within each household 10,000 times, and recalculated the effect size for Native American ancestry in each round, to make an empirical null distribution. None of the permutations resulted in a greater effect size than that of the original data (permutation effect size ranging from -5.6 cm to 5.8 cm, permutation mean effect size = 0 cm, observed effect size = -14.7 cm,) suggesting that our height association could not be explained by a household effect.

**Polygenic risk score analysis**

Previous large-scale height GWAS, done predominantly in Europeans, have identified 3,290 independent common height-associated variants (*21*). To assess the predictive power of these European-biased variants in the Peruvian population we generated polygenic risk scores (PRS) based on the reported effect sizes of 2,993 common height-associated variants that were present in our cohort. Out of these variants, 1,519 (51%) showed directionally consistent effects between our Peruvian GWAS and the European GWAS (*21*), and 199 (7%) had p-value < 0.05 in our Peruvian GWAS. Higher PRS bins were associated with increased height ($r$ = 0.2, p-value = $2.7 \times 10^{-34}$). The estimated genetic heritability ($h_g^2$) of height was similar for Peruvians ($h_g^2$ = 57.6%) and Europeans ($h_g^2 = 62.5\%$) (*30*); however, previously identified height-associated variants explained only 6.1% of height phenotypic variance in our cohort compared to 24.6% in the original European cohort**,** suggesting that either different variants are responsible for the height variance in the Peruvian population or the lead European variants do not tag the same causal variants in the Peruvian population. This observation is in line with a number of recent reports (*16*, *31*, *32*) showing the lower predictive power of PRS calculated based on European

GWAS in non-European populations as a result of differences in demographic history and linkage disequilibrium (LD) patterns.

**Functional annotation of rs200342067**

Commonly used variant annotation tools predict rs200342067 to have a severe functional consequence (scaled Combined Annotation Dependent Depletion (CADD) score (*33*): 33 (e.g. top 0.1% of all single nucleotide changes), SIFT prediction (*34*): "deleterious", PolyPhen prediction (*35*): "probably damaging"). These predictions, although unlikely in light of our findings, are expected since rs200342067 is extremely rare or absent in most human populations (MAF = 0.1%, Genome Aggregation Database (gnomAD), N = 141,456) (*36*), and other missense mutations in *FBN1* are known to cause nine different Mendelian diseases, all of which are dominantly inherited (*37*).

**Genomic context of rs200342067**

The rs200342067 variant changes the conserved T (ancestral) allele to a C (derived) allele in *FBN1* exon 31 (g.48773926T>C). This change substitutes a glutamic acid, a large amino acid with a negatively charged side chain, in position 1,297 with a glycine, the smallest amino acid with no side chain, (p.Glu1297Gly). p.Glu1297Gly is located in Fibrillin-1 calcium binding epidermal growth factor domain 17 (cbEGF-domain 17), between a conserved cysteine (p.Cys1296) involved in forming a disulfide bond with p.Cys1284, and a conserved asparagine (p.Asp1298) involved in calcium binding (*29*).

**Supplementary Figures**

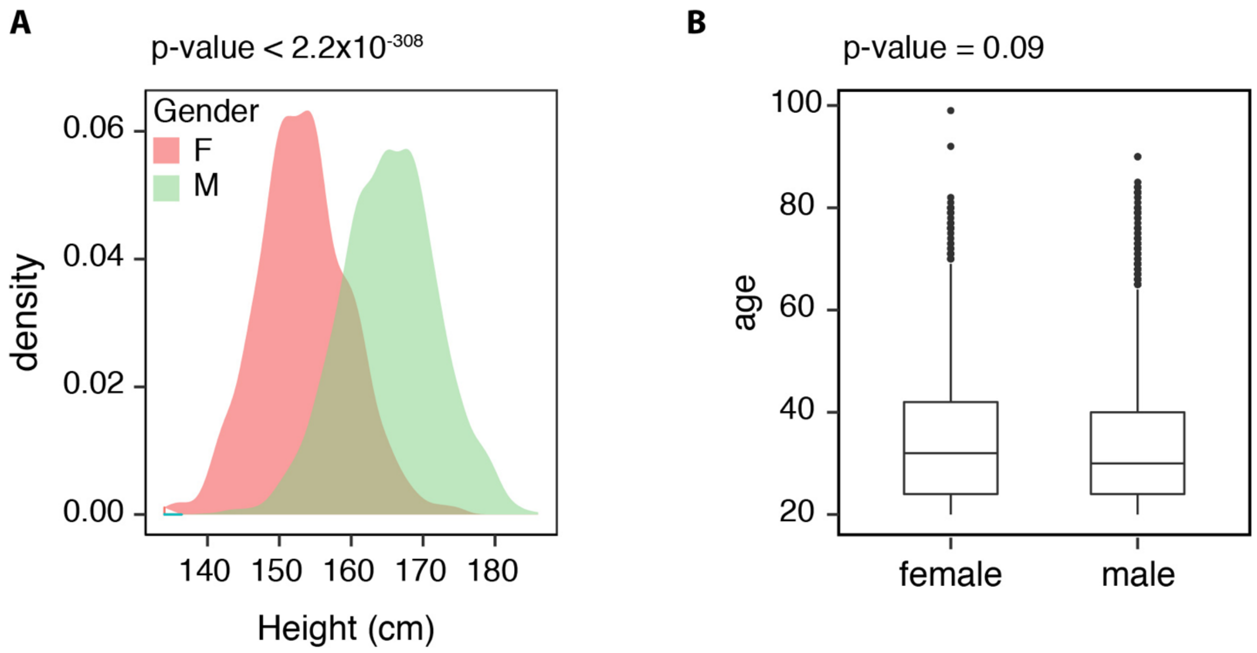


**Figure S1**: **Cohort's demographic information. A)** Density plot of height for all the Peruvian males (N = 1,795 (57%)) and females (N = 1,339 (43%)) included in this study after quality control (e.g. after removing low quality samples, individuals below 18 years old and height outliers (± 3 x standard deviations (SD) from the mean). Males were significantly taller than females (Male mean = 165.2 cm (SD = 6.7), Female mean = 153.4 cm (SD = 6.4), p-value < $2.2x10^{-308}$). **B**) Age was not significantly different between males and females (t-test p-value = 0.09).

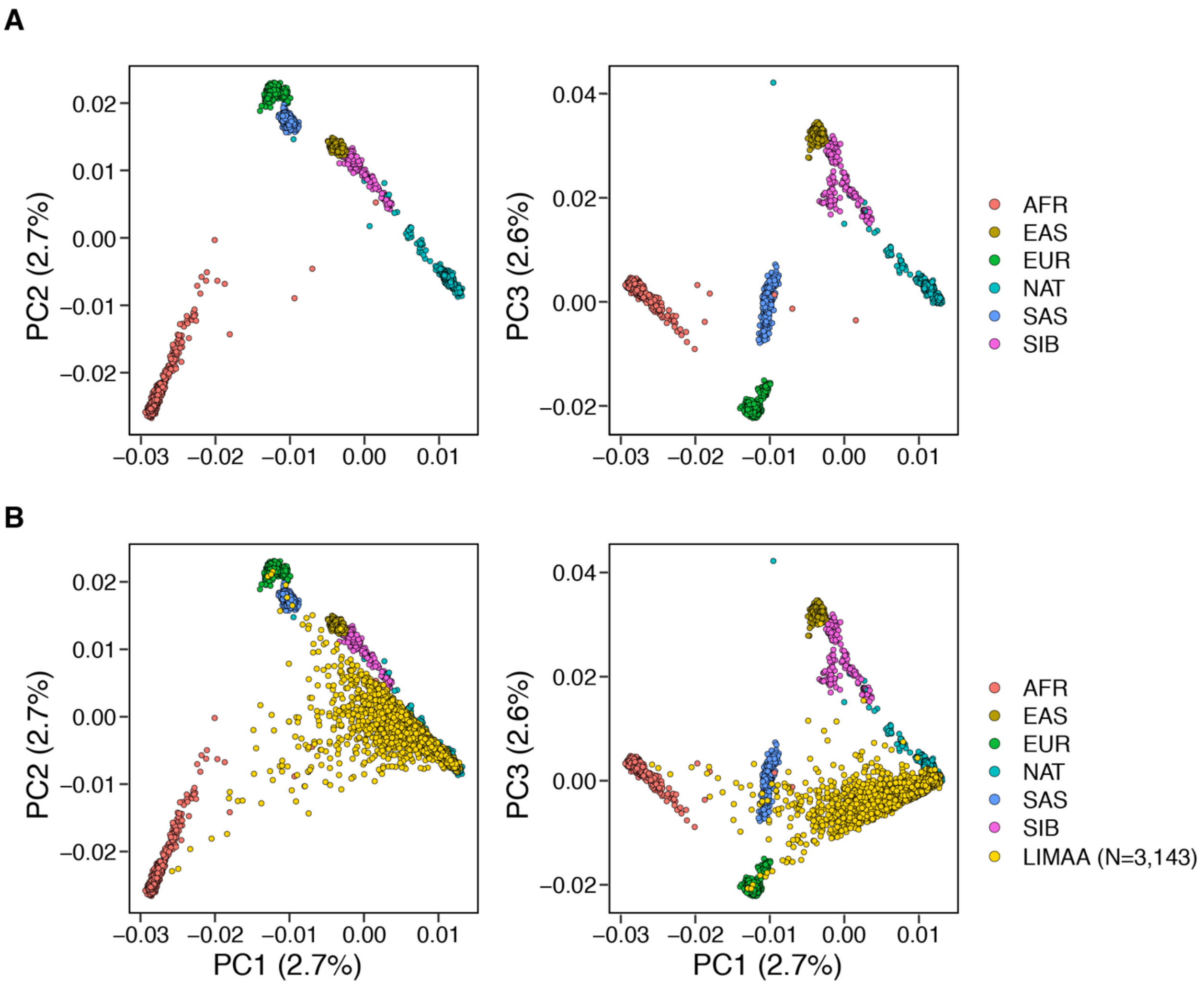


**Figure S2: Principal component analysis (PCA).** PCA analysis of genotyping data from Peruvians included in this study merged with the data from continental populations from the 1000 Genomes Project phase 3 (N = 3469) (*1*, *2*) as well as the data from Siberian and Native American populations from Reich et *al*. 2012 *Nature* study (*3*) (N = 738) as reference panel (number of variants = 34,936, MAF > 1%, genotype missingness < 5%). In order to better visualize the relative position of reference populations we plotted the data **A)** without and **B)** with the Peruvians from this study (N = 3,134)**.** Each individual is represented as a dot. Populations are colored based on their continental origin for the 1000 Genomes Project data and based on assignment to Native American or Siberian tribes for the Reich data (AFR: Africa, AMR: South America, EAS: East Asia, SAS: South Asia, EUR: Europe, SIB: Siberian, NAT: Native American).

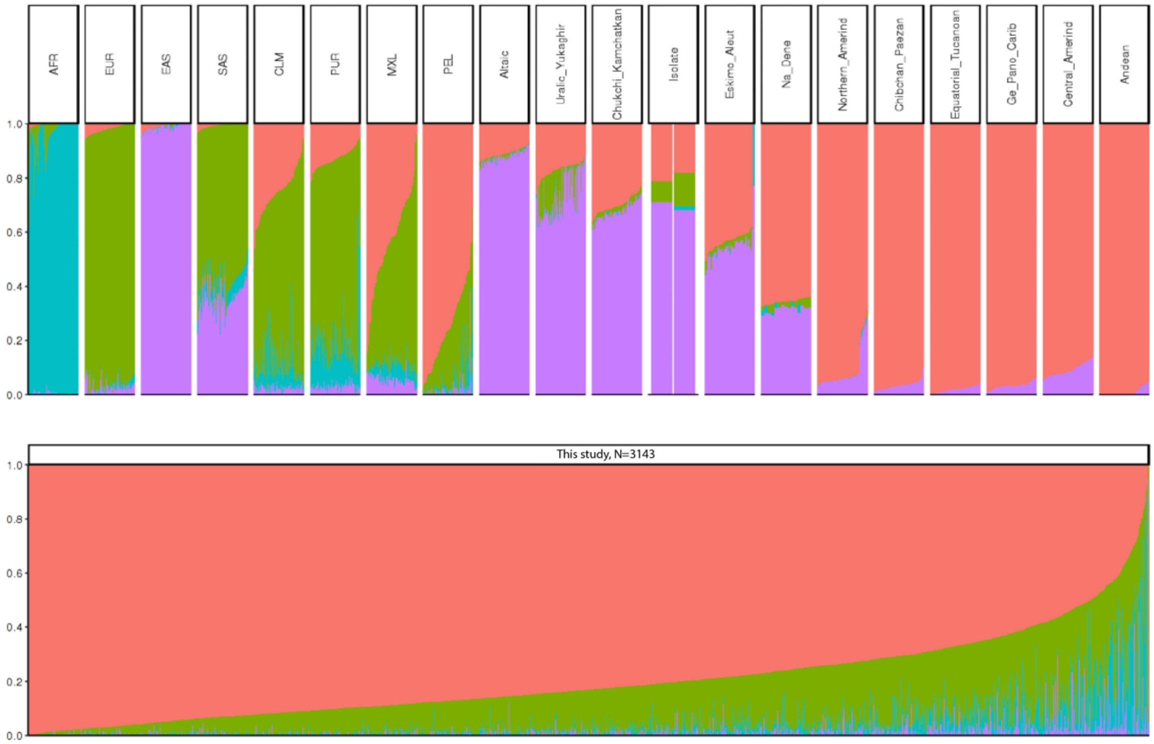


**Figure S3: Global ancestry analysis using ADMIXTURE (K=4)**. We observed varying levels of European, African, and Asian admixture in the Peruvian population with a median proportion of Native American, European, African, and Asian ancestry per individual of 0.83 (Interquartile range (IQR) = 0.72-.91), 0.14 (0.08-0.21), 0.01 (0.003-0.03), and 0.003 ($10^{-5}$-0.01) respectively. Each individual is represented as a thin vertical line, each color corresponds to the genomic proportion of a given cluster in that individual's genome. The ADMIXTURE with K=4 analysis is done using all populations in 1000 Genomes Project phase 3 (*1*, *2*) and Siberian and Native American populations from the Reich et *al*. 2012 *Nature* study (*3*). AFR: African ancestry includes :Yoruba in Ibadan, Nigeria, Luhya in Webuye, Kenya, Gambian in Western Divisions in the Gambia, Mende in Sierra Leone, Esan in Nigeria, Americans of African Ancestry in SW USA; EUR: European ancestry, includes: Central European, Utah Residents (CEPH) with Northern and Western European Ancestry, Toscani in Italy, Finnish in Finland, British in England and Scotland, Iberian Population in Spain; EAS: East Asian, includes: Han Chinese in

Beijing, China, Japanese in Tokyo, Japan, Southern Han Chinese, Chinese Dai in Xishuangbanna, China, Kinh in Ho Chi Minh City, Vietnam; SAS: South Asian, includes: Gujarati Indian from Houston, Texas, Punjabi from Lahore, Pakistan, Bengali from Bangladesh, Sri Lankan Tamil from the UK, Indian Telugu from the UK; PUR: Puerto Ricans from Puerto Rico; CLM: Colombian from Medellin, Colombia; MXL: Mexicans from Los Angeles, California; PEL: Peruvians from Lima, Peru. Altic: Altaic language family, includes: Yakut, Buryat, Evenki, Tuvinians, Altaian, Mongolian, Dolgan. North Amerind: Northern Amerindian language family, includes: Maya, Mixe, Kaqchikel, Algonquin, Ojibwa, and Cree. Central Amerind: Central Amerindian language family, includes: Pima, Chorotega, Tepehuano, Zapotec, Mixtec, and Yaqui. Andean: Andean language family, includes: Quechua, Aymara, Inga, Chilote, Diaguita, Chono, Hulliche, and Yaghan. For a full list of all populations in all language groups please see the Reich et *al*. 2012 *Nature* study (*3*)**.**

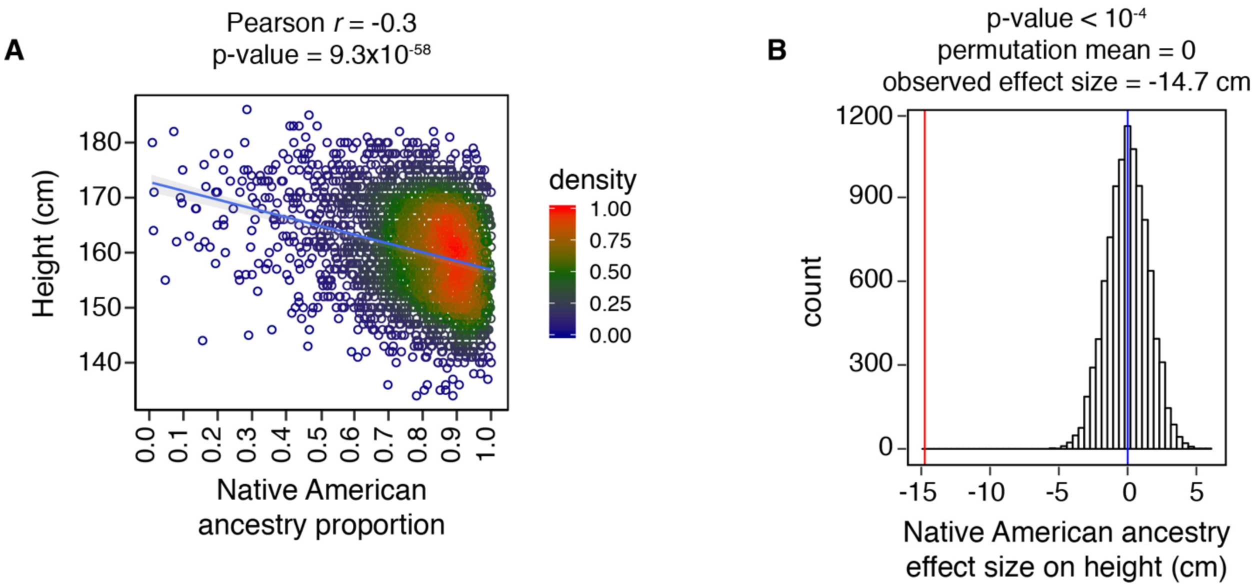


**Figure S4: The effect of Native American ancestry proportion on height. A)** Greater Native American ancestry proportion is associated with lower height (N=3,134, $r$ = -0.3, p-value = $9.3 \times 10^{-58}$). The x-axis represents Native American ancestry proportion from ADMIXTURE analysis at K = 4 clusters. The y-axis represents height (cm). **B)** Height was randomly reassigned to individuals within each household, and the effect size of Native American ancestry on height was recalculated to derive an empirical null distribution of effect sizes. None of the permutations resulted in a greater effect size than that of the original data (permutation effect size ranging from -5.6 cm to 5.8 cm, permutation mean effect size = 0 cm, observed effect size = -14.7 cm)

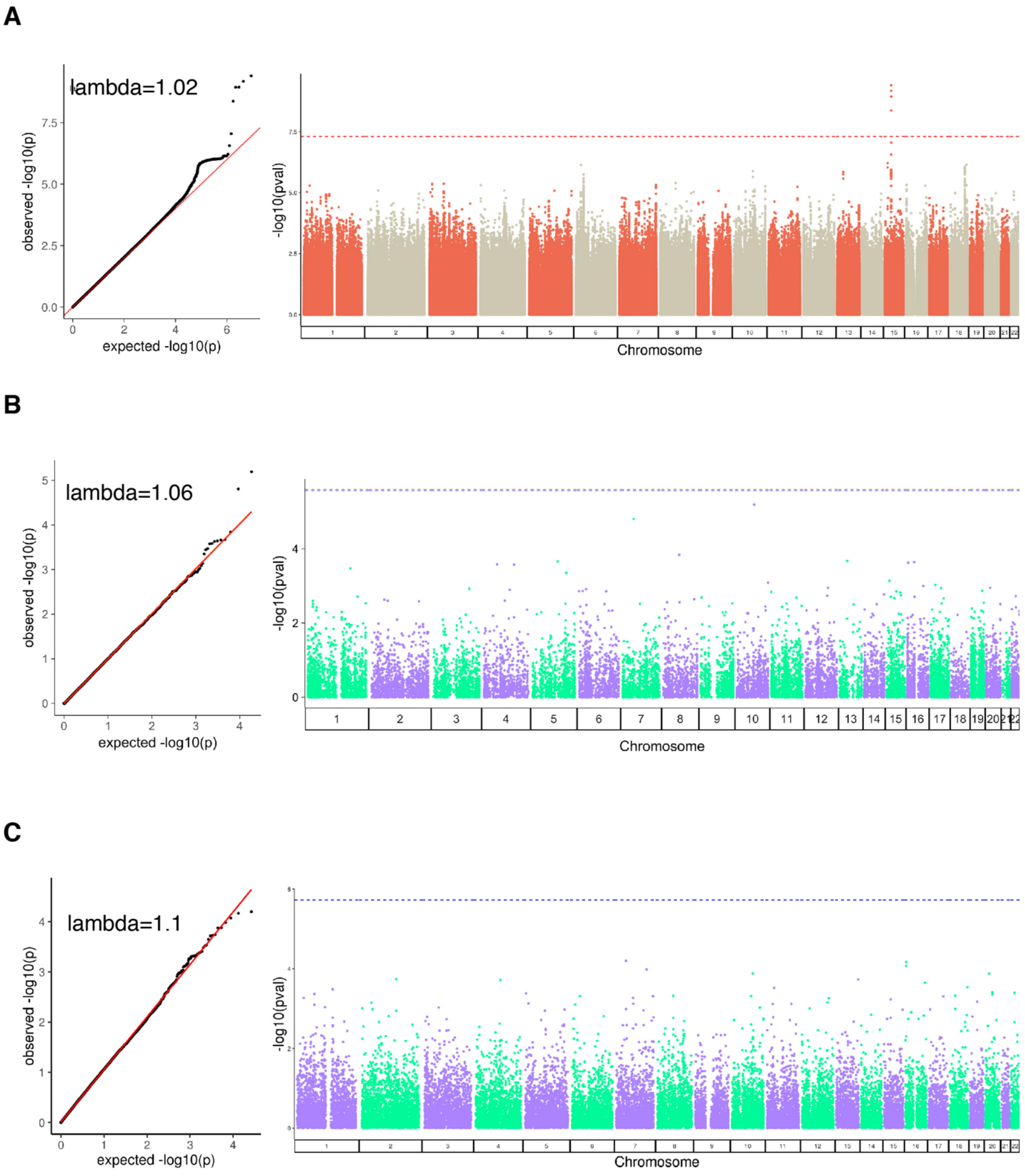


**Figure S5: Manhattan and quantile-quantile (QQ) plots. A)** Single variant association analysis using GEMMA (*4*), the dotted red line corresponds to the genome-wide significance threshold of $5x10^{-8}$ for single variant association testing. Five SNPs passed the genome-wide significance threshold. **B)** Rare (MAF < 1%) variants gene-based analysis using SKAT (*5*) the dotted red line corresponds to the genome-wide significance threshold of $2x10^{-6}$ for 25,000 tested genes. No SNPs reached the genome-wide significance threshold. **C)** gene-based meta-analysis of common variants using GCTA fastBAT (*6*) the dotted red line corresponds to the genome-wide significance threshold of $2x10^{-6}$ for 25,000 tested genes. No SNPs reached the genome-wide significance threshold.

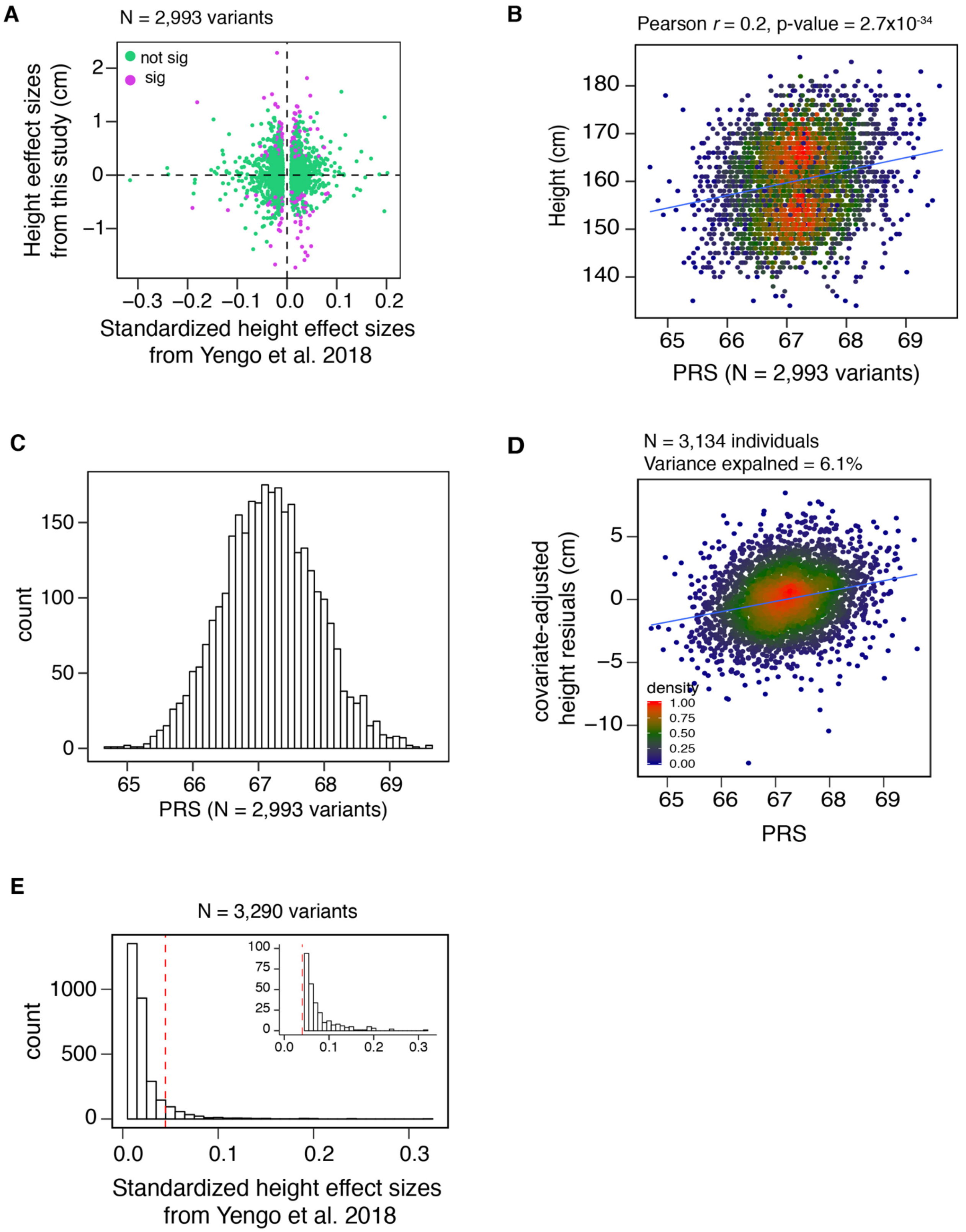
A
N = 2,993 variants
not sig
sig
Height eeffect sizes from this study (cm)
Standardized height effect sizes from Yengo et al. 2018
B
Pearson r = 0.2, p-value = 2.7x10-34
Height (cm)
PRS (N = 2,993 variants)
C
count
PRS (N = 2,993 variants)
D
N = 3,134 individuals
Variance expalned = 6.1%
covariate-adjusted height resiuals (cm)
density
PRS
E
N = 3,290 variants
count
Standardized height effect sizes from Yengo et al. 2018

**Figure S6: Polygenic risk score (PRS) analysis.** We used effect sizes from 2,993 common height-associated variants from the Yengo et al 2018 meta-analysis (N ~ 700,000 European individuals) (*7*) that were present in our cohort (N = 3,134 Peruvian individuals) to derive the PRS. **A)** Out of 2,993 variants, 1,519 (51%) showed directionally consistent effects, and 199 (7%) had p-value < 0.05 in our Peruvian GWAS. **B)** Higher PRS values are associated with increased height ($r$ = 0.2, p-value = $1.7x10^{-34}$). **C)** Histogram showing the PRS distribution. **D)** Previously identified height-associated variants explained only 6.1% of height phenotypic variance in our cohort ($r$ = 0.061, p-value = $6.8x10^{-45}$) , x-axis: PRS, y-axis: height residuals after adjustments for age and gender as fixed effects and a GRM as random effect. **E)** The majority (99%) of previously identified common height-associated variants (N = 3,290) have effects less than 5 mm per allele (dashed red line: cutoff corresponding to 5 mm effect size, smaller plot shows the zoomed in tail of the main plot).

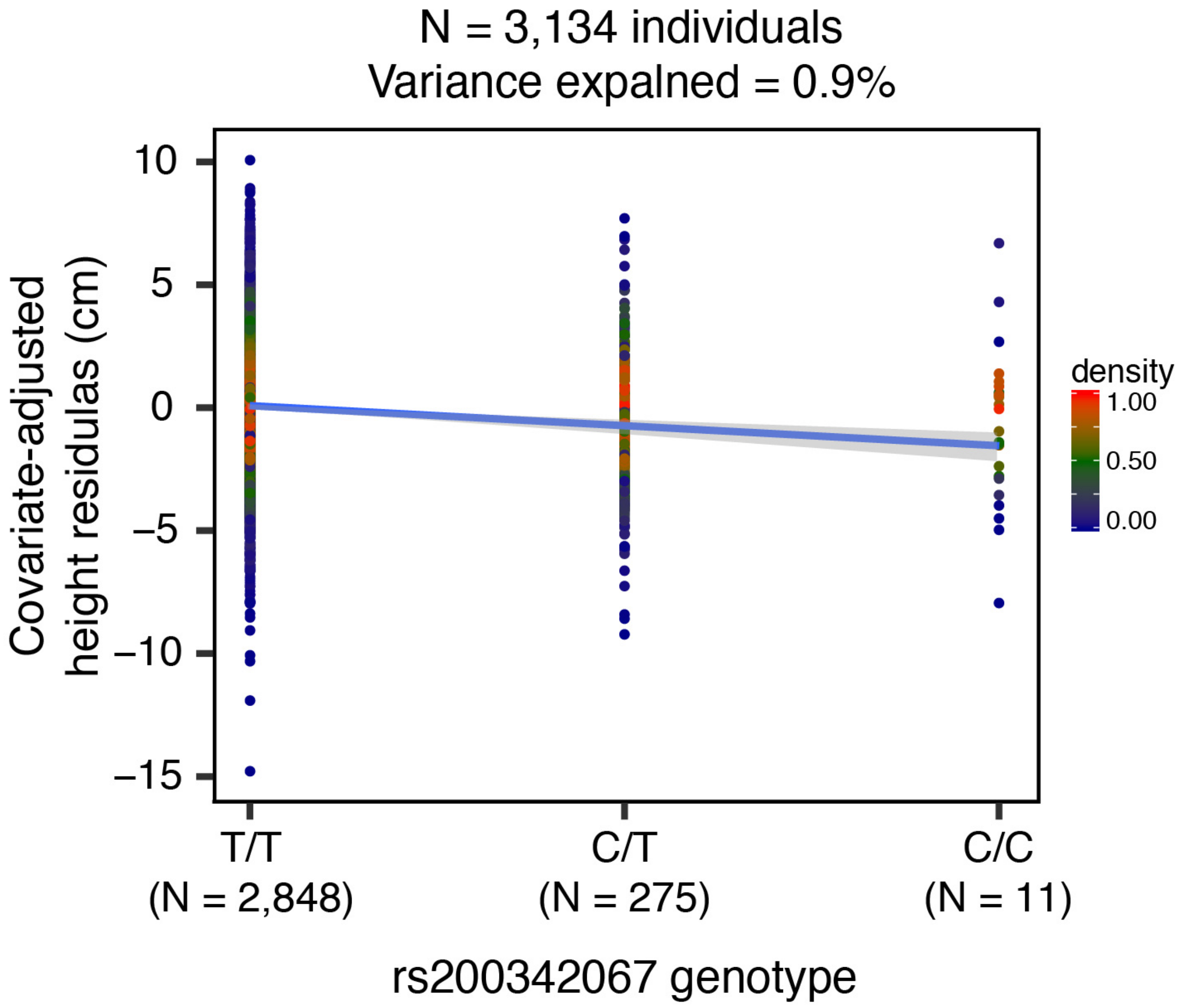


**Figure S7: Effect size of rs200342067 on height in the Peruvian population**. rs200342067 in heterozygous individuals reduces height by 2.2 cm (4.4 cm in homozygous individuals, including 11 individuals with C/C genotype, 275 C/T genotype, and 2,848 T/T genotype) and could explain 0.9% of height phenotypic variance in our cohort (N = 3,143). x-axis: rs200342067 genotype, y-axis: height residuals after adjustments for age and gender as fixed effects and a GRM as random effect.

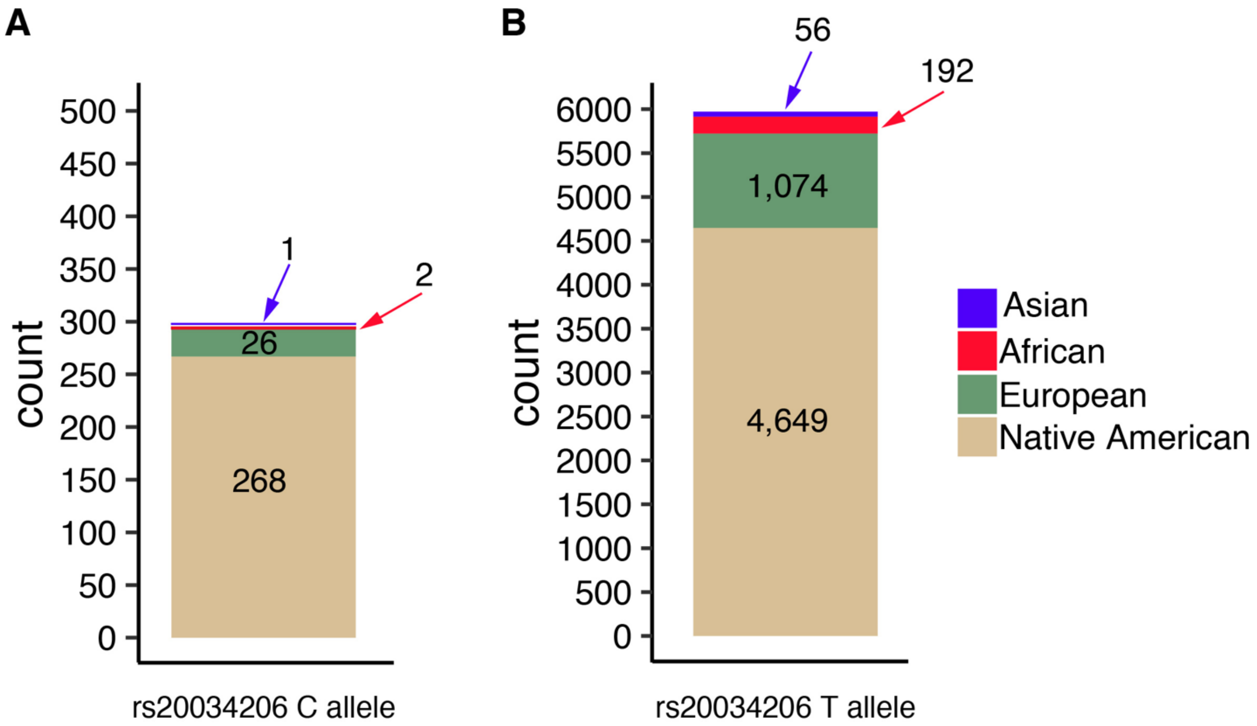


**Figure S8: Local ancestry inference at the rs20034206 locus.** To test for positive selection at the rs20034206 locus, we restricted the analysis to haplotypes in which the local ancestries of both C and T alleles were inferred to be Native American. **A)** Local ancestry inference results for A) rs20034206 C allele, and **B)** rs20034206 T allele.

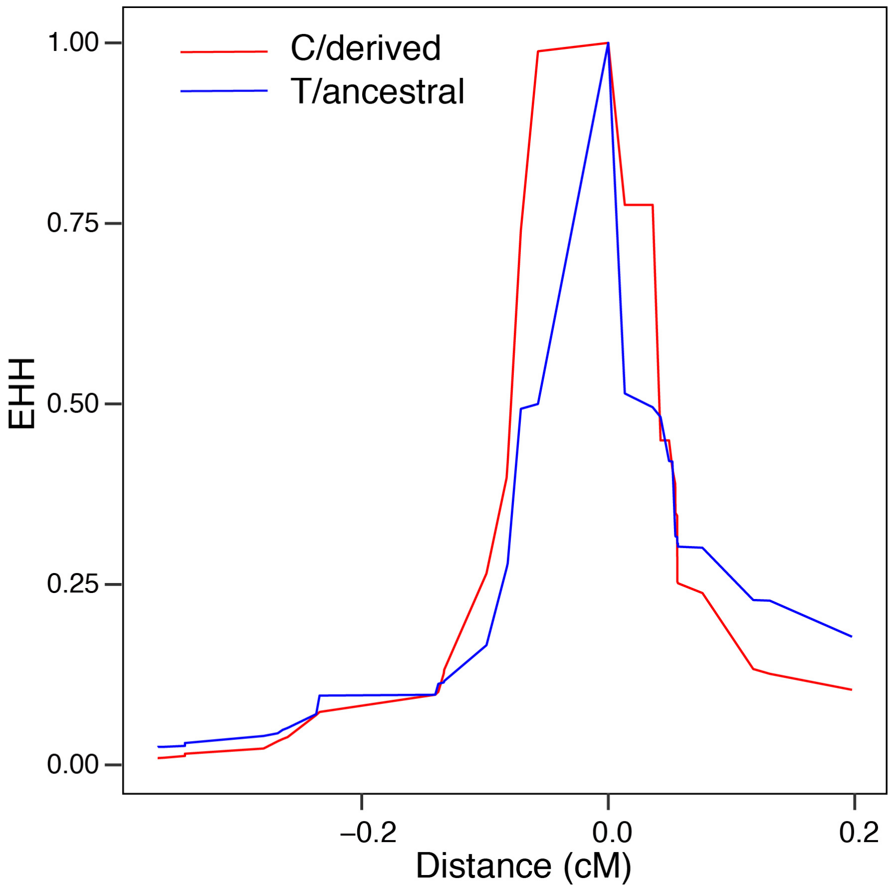


**Figure S9: Extended haplotype homozygosity (EHH) for rs20034206, C and T alleles.** Haplotypes carrying the C allele show a slower decay of homozygosity compared to the haplotypes carrying the T allele. Analysis is restricted to haplotypes in which rs20034206 is inferred as Native American.

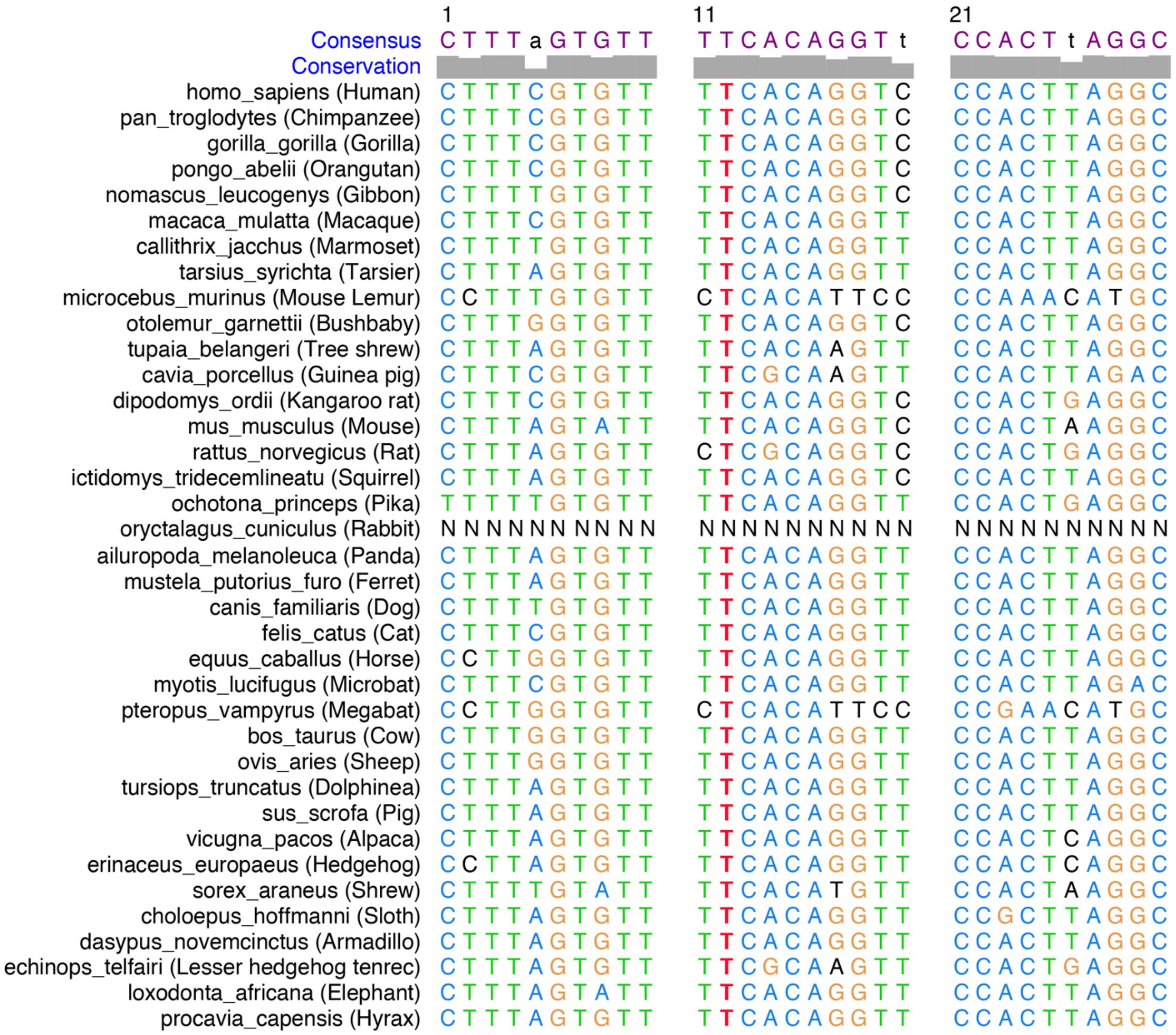


**Figure S10: Multiple sequence alignment around rs20034206 in 37 eutherian mammals**. rs200342067 changes a conserved T allele (ancestral, shown in red) to a C allele (derived). Sequence alignments were obtained from Ensembl GRCh37 release 95.

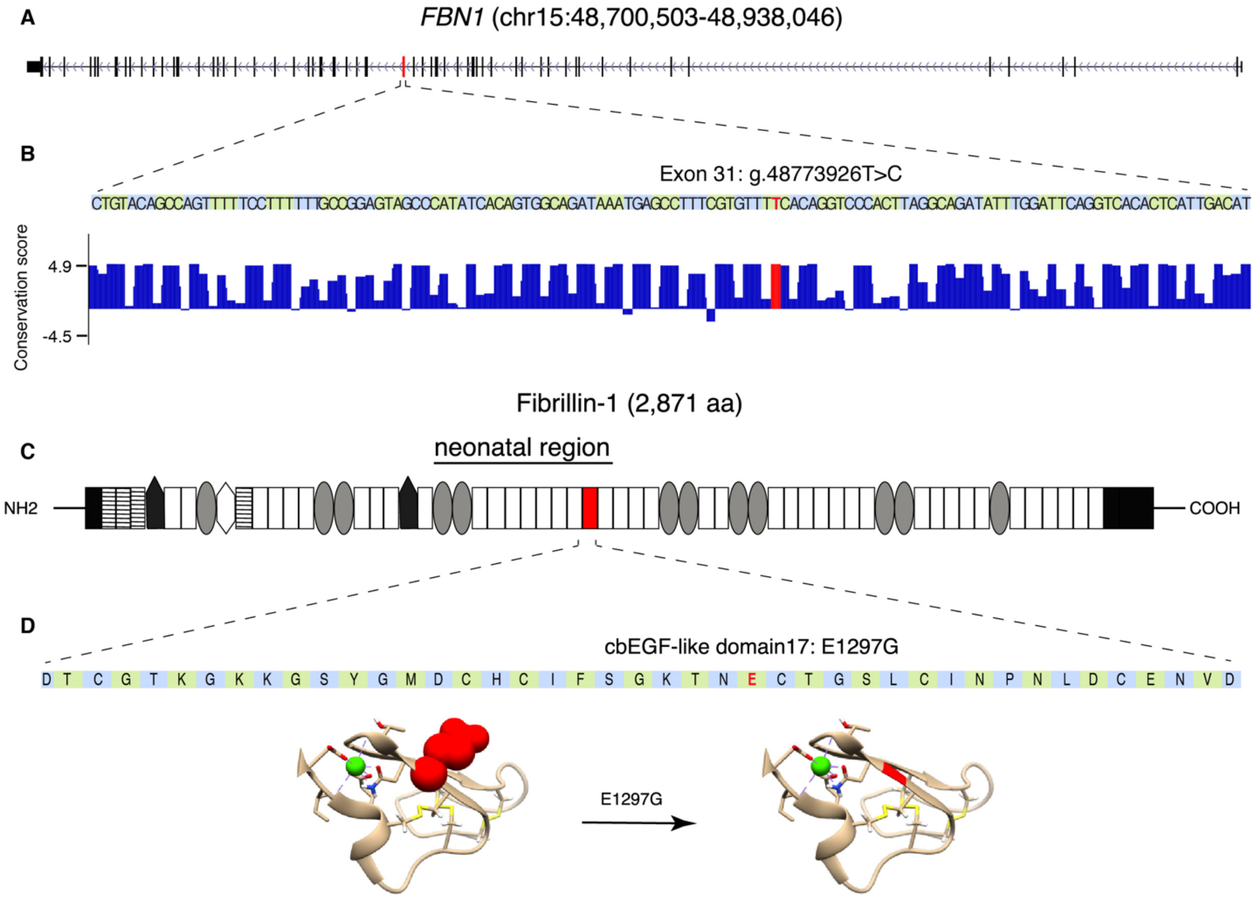


**Figure S11: Genomic context of rs200342067 (p.Glu1297Gly). A)** Schematic representation of *FBN1*, exons are shown as black bars. Exon 31 (ENSE00001753582) is shown in red. B) *FBN1* exon 31 sequence and PhyloP per-nucleotide conservation score based on multiple alignment of 100 vertebrate species (obtained from UCSC genome browser GRCh37 assembly, conservation track). The T>C change due to rs200342067 occurs in a conserved nucleotide. **C)** Schematic representation of Fibrillin-1 (ENST00000316623.5). Fibrillin-1 consists of: N and C terminal (black rectangles), EGF-like domains (stripped rectangles), hybrid domains (black pentagons), TGFβ-binding domains (gray ovals), a proline-rich domain (white hexagon), and 43 calcium binding cbEGF-like domains (white rectangles). cbEGF-domain 17, the domain affected by rs200342067 (p.Glu1297Gly), is shown in red. p.Glu1297Gly is located between a conserved cysteine (p.Cys1296) involved in forming a disulfide bond with p.Cys1284 and a conserved asparagine (p.Asp1298) involved in calcium binding. **D)** Fibrillin-1 cbEGF-domain 17 sequence and 3D structure of cbEGF-domains 17 and 18 (the 3D structure was obtained based homology with fibrillin-1 cbEGF-domains 12 and 13, 1LMJ (*8*), in the Protein Data Bank). rs200342067 changes glutamic acid, a large amino acid with a negatively charged side chain, to glycine, the smallest amino acid with no side chain (shown in red). The side chains are shown for rs200342067 (red spheres), the calcium-interacting residues (beige sticks), and the cysteine residues involved in disulfide bonds (yellow sticks). Calcium ion is shown in green.

**Table S1: Base Model parameters.** Native American ancestry is significantly associated with lower height after accounting for age, gender, African and Asian ancestry proportions, and a genetic relatedness matrix (GRM) to account for population structure and genetic relatedness. ASI: Asian, AFR: African, EUR: European, NAT: Native American.

| term | estimate | std.error | statistic | Pr.Chi. (df=1) | 2.50% | 97.50% |
|---|---|---|---|---|---|---|
| age | -0.10 | 0.01 | -12.96 | 1.50E-37 | -0.12 | -0.09 |
| Gender M | 11.33 | 0.22 | 50.57 | 0.00E+00 | 10.90 | 11.77 |
| AFR proportion | -3.25 | 2.15 | -1.51 | 1.31E-01 | -7.46 | 0.97 |
| ASI proportion | -10.73 | 3.38 | -3.18 | 1.49E-03 | -17.34 | -4.11 |
| NAT proportion | -14.55 | 1.04 | -14.00 | 2.43E-43 | -16.59 | -12.52 |

**Table S2: Base model plus a household random effect.** Native American ancestry remained significantly associated with lower height after we included a random household effect as a proxy for socioeconomic and environmental factors.

| term | estimate | std.error | statistic | Pr.Chi. (df=1) | 2.50% | 97.50% |
|---|---|---|---|---|---|---|
| age | -0.10 | 0.01 | -12.43 | 1.06E-34 | -0.12 | -0.09 |
| Gender M | 11.47 | 0.22 | 51.20 | 0.00E+00 | 11.03 | 11.91 |
| AFR proportion | -3.57 | 2.15 | -1.66 | 9.63E-02 | -7.77 | 0.64 |
| ASI proportion | -11.62 | 3.40 | -3.42 | 6.38E-04 | -18.28 | -4.95 |
| NAT proportion | -14.75 | 1.06 | -13.94 | 7.20E-43 | -16.83 | -12.68 |
| sd_(Intercept).household | 2.08 | NA | NA | 7.40E-07 | 1.54 | 2.53 |

**Table S3: Association signal at 15q15-21.1**. This locus overlaps the coding sequence of *FBN1* and includes five single nucleotide polymorphisms (SNPs), which are in high LD.

| chromosome | rs ID | position | allele1 | allele0 | allele1 frequency | effect size (cm) | se | p-value |
|---|---|---|---|---|---|---|---|---|
| 15 | rs193211234 | 48752674 | A | T | 0.046 | -2.4 | 3.79E-01 | 4.90E-10 |
| 15 | rs200342067 | 48773926 | C | T | 0.047 | -2.2 | 3.59E-01 | 7.96E-10 |
| 15 | rs544786245 | 48822780 | T | G | 0.044 | -2.3 | 3.79E-01 | 1.26E-09 |
| 15 | rs143730951 | 48858921 | T | C | 0.047 | -2.3 | 3.71E-01 | 1.29E-09 |
| 15 | rs180913076 | 48928052 | C | A | 0.045 | -2.2 | 3.77E-01 | 4.71E-09 |

**Table S4: Inclusion of other covariates in rs200342067 association testing.** Inclusion of principal components (PCs), socioeconomic status (SES), or ancestry proportions (ASI: Asian, AFR: African, EUR: European) did not change the association effect size or strength.

| Covariates | rs200342067 effect size (cm) | SE | rs200342067 association p-value (score test) |
|---|---|---|---|
| Age, gender, GRM | -2.2 | 4E-01 | 8.0E-10 |
| Age, gender, 10 PCs, GRM | -2.2 | 4E-01 | 9.5E-10 |
| Age, gender, 10 PCs, SES, GRM | -2.2 | 4E-01 | 1.5E-09 |
| Age, gender, 20 PCs, GRM | -2.2 | 4E-01 | 3.0E-09 |
| Age, gender, ASI, AFR, EUR, GRM | -2.2 | 4E-01 | 9.5E-10 |

**Table S5: rs200342067, C genotype carriers in *BioMe* cohort**. Individuals are stratified by country of origin. No homozygous individual (C/C) was observed in *BioMe.* For the replication analysis, we restricted the age to ≥ 18 and ≤ 80 for females, and ≥ 22 and ≤ 80 for males.

| Country | N Carriers | N Total | MAF (%) |
|---|---|---|---|
| Argentina | 1 | 63 | 0.793651 |
| Cuba | 3 | 163 | 0.920245 |
| Ecuador | 10 | 431 | 1.160093 |
| Guatemala | 2 | 71 | 1.408451 |
| Mexico | 5 | 318 | 0.786164 |
| Peru | 9 | 124 | 3.629032 |
| USA | 8 | 20877 | 0.01916 |

**Table S6: Comparison of rs200342067 minor allele count between populations from different geographical regions in Peru.** rs200342067 was significantly more frequent in Coastal populations than in populations from the Andes and the Amazon.

| Geographical region | A1 | A2 | MAC | # individuals | observed MAF (%) |
|---|---|---|---|---|---|
| **andes** | C | T | 2 | 56 | 1.7 |
| **amazon** | C | T | 0 | 23 | 0 |
| **coast** | C | T | 7 | 36 | 9.7 |
| **All** | **C** | **T** | **9** | **116** | **3.9** |

**Table S7: Disease, phenotypes, and traits caused by mutations in FBN1.**

| Disease | OMIM ID | Inheritance |
|---|---|---|
| Acromicric dysplasia | 102370 | AD |
| Ectopia lentis | 129600 | AD |
| Geleophysic dysplasia | 614185 | AD |
| Marfan lipodystrophy syndrome | 616914 | AD |
| Marfan syndrome | 154700 | AD |
| MASS syndrome | 604308 | AD |
| Stiff skin syndrome | 184900 | AD |
| Weill-Marchesani syndrome | 608328 | AD |
| Shprintzen-Goldberg craniosynostosis syndrome | 182212 | AD |

**Table S8: Demographic information of clinical examination participants.** Skin biopsies were obtained from 11 including: 2 with C/C, 2 with C/T, and 7 with T/T genotypes at rs200342067.

| | Age | Gender | Height cm | EUR | NAT | AFR | ASI | C allele count at rs200342067 |
|---|---|---|---|---|---|---|---|---|
| Individual 1 | 64 | F | 146 | 0.020 | 0.003 | 0.975 | 0.002 | 2 |
| Individual 2 | 35 | F | 144 | 0.213 | 0.000 | 0.782 | 0.005 | 2 |
| Individual 3 | 30 | F | 146 | 0.052 | 0.003 | 0.932 | 0.013 | 1 |
| Individual 4 | 60 | M | 164 | 0.166 | 0.024 | 0.810 | 0.001 | 1 |
| Individual 5 | 56 | M | 164 | 0.125 | 0.003 | 0.865 | 0.008 | 0 |
| Individual 6 | 37 | F | 160 | 0.201 | 0.016 | 0.782 | 0.001 | 0 |
| Individual 7 | 30 | F | 167 | 0.072 | 0.000 | 0.928 | 0.000 | 0 |
| Individual 8 | 60 | F | 157 | 0.047 | 0.000 | 0.953 | 0.000 | 0 |
| Individual 9 | 46 | F | 153 | 0.071 | 0.000 | 0.929 | 0.000 | 0 |
| Individual 10 | 44 | F | 150 | 0.032 | 0.002 | 0.952 | 0.015 | 0 |
| Individual 11 | 36 | F | 154 | 0.087 | 0.003 | 0.678 | 0.232 | 0 |
